# Activity in the peripheral representation within primate V1 is substantially modulated during running

**DOI:** 10.1101/2024.10.10.617723

**Authors:** Declan P. Rowley, Alexander C. Huk, Jacob L. Yates

## Abstract

We recently investigated whether activity in primary visual cortex of a primate (Callithrix jacchus) is modulated during running, and found that the effects were small (and suppressive), a notable difference from the large and positive modulations observed in mice. In that first report, we noted that the majority of our data were collected from the retinotopic representation of the fovea, and surmised that running modulations might be different in the peripheral representation. Here, we report that running-correlated modulations of the peripheral representation in marmoset V1 are positive and substantial— on order of 30%. In light of both the small and negative modulations observed in foveal V1, and the large and positive modulations seen in mouse V1, these results suggest that the foveal representation in primates may be unique. In this domain, non-foveal V1 in primates appears more similar to that of rodents.

## Introduction

Although activity in mammalian primary visual cortex (V1) is known to receive strong inputs that originate in the retina, the degree to which nonvisual factors affect its processing has remained a topic of serious investigation [1–3]. Based on studies of attention-related modulations in primate V1, a broadly-held assumption in the field was that the behavioral / attentive state of the organism exerted modest, if any, effects on visual processing in this first cortical stage. However, studies in mice showed that when animals ran, activity in their V1 was strongly modulated [4]. In a recent report, we attempted to resolve this tension between species by performing matched running-modulation tests in marmosets, a small new world primate amenable neurophysiology and eye tracking while perched on a treadmill apparatus. We found that marmoset V1 was not much affected by running, although we did detect (via three different types of analysis) a small suppressive modulation during running. Thus, while both mouse and marmoset V1 demonstrated running-correlated changes (both of which could be descried well by a shared gain term that was correlated with running), the smaller magnitude and opposite sign of the primate V1 modulations suggested a severe species difference in how robust V1 is to nonvisual factors.

In that first report, we relied on data from 2 types of electrophysiological recording device; we started the experiments using chronically-implanted N-form arrays, but later developed techniques that allowed us to use Neuropixels 1.0 (mouse-style) probes in the daily-acute recording style [5]. Because the foveal representation in marmosets is located on the the superficial surface of the dorsal occipital lobe, our chronically-implanted arrays (which produced the majority of that first dataset) sampled neurons in this region. Our later Neuropixels recordings sampled both the foveal representation and (by virtue of hitting deeper folds of the occipital lobe, such as the calcarine sulcus) more peripheral portions of V1’s retinotopic map. In those few peripheral recordings, we noted that the running modulations were different than in the foveal representation. Given that hint, we decided to refine and focus our Neuropixels recording approach to collect a larger and more definitive dataset, allowing us to more comphrehensively compare running modulations from simultaneously-recorded portions of V1 representing multiple regions of the retinotopic map. Here, we report these results, meant to be a more definitive follow-up to that prior report. We found that running-correlated modulations in peripheral V1 were positive and much larger than the subtle effects we previously observed in the foveal representation.

Before reporting our results in detail, it is important to clarify our working definition of the fovea (and by exclusion, of the periphery). We ground our definition in the retina, where the fovea is a central portion with well-known anatomical specialization. The size of the fovea in marmosets corresponds to the central 8-10 deg of visual space (and is similar in both anatomical and physiological properties across anthropoids [6, 7]). The specialized anatomical and functional aspects of this region of the retina might thus be expected to carry forward with corresponding functional uniqueness in the cortical representations of the central 10 deg of the visual field. Due to looseness of terminology in the cortical and psychophysical literature, here we emphasize that the fovea should be distinguished from the ‘foveala’, the highest acuity region within the fovea (on order of a single degree of visual angle). Although that highest-acuity central degree of vision is certainly functionally interesting and important, here we focus on the larger anatomical fovea and the corresponding retinotopic representations in cortex.

## Results

We measured V1 activity in head-fixed marmosets on a wheel-based treadmill. Animals were free to run (or to perch without running) while they freely viewed visual stimuli on a computer monitor. We recorded neural activity using Neuropixels 1.0 probes with recording sites located within the foveal representation as well as in other more eccentric portions of the retinotopic map. Our methods mirror our previous report [3], with the modification that we focus exclusively on datasets in which we simultaneously recorded from foveal and peripheral sites in V1 (Figure 1).

**Fig. 1:**
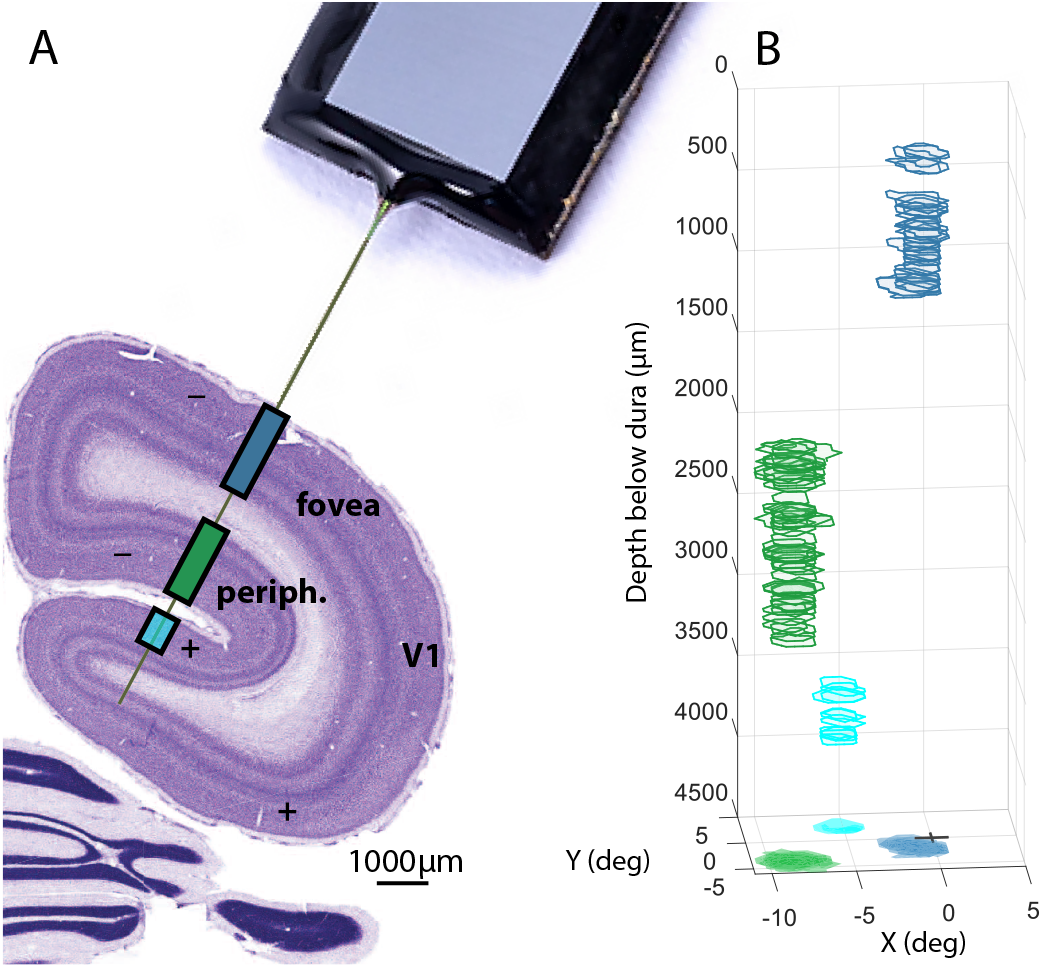
Simultaneous foveal and peripheral columnar recording paradigm. A. The marmoset visual cortex is folded such that the peripheral representation is within the calcarine sulcus. Using Neuropixels probes we were able to record from multiple banks of V1 simultaneously. The foveal representation of the lower field (purple) is immediately below dura the burr hole on the dorsal surface of the cortex on the most proximal part of the probe. Beneath this, after the probe passes through some white matter it reaches a second, but more peripheral, lower field representation (green). In some cases, the probe reached a third peripheral, this time upper field, representation (cyan). Coronal Nissl stain from Paxinos et al. (2008) at 8.5mm posterior of ear bar zero. B. 50% contours of receptive fields, obtained from reverse correlation of free viewing of sparse dots and fit with a gaussian, by depth on probe, with banks in colors corresponding to A. The x-y plane at the bottom of the figure includes a projection of all the receptive fields to demonstrate the high amount of overlap within banks of these columnar recordings.

At the beginning of each session, we mapped the receptive fields of the recorded neurons using techniques previously described and used for free-viewing marmosets [8]. As expected, neurons in superficial recording sites had receptive fields within the central few degrees of the visual field (corresponding to the foveal representation); neurons encountered in deeper portions of V1 (i.e., in the calcarine sulcus) had receptive fields that spanned more eccentric locations (with angular locations that were consistent with the known retinotopic map of marmoset V1). These V1 neurons also exhibited standard forms and degrees of orientation selectivity. These basic observations confirm that the neurons we recorded from could be considered standard primate V1 neurons, well-driven by classical stimuli and with standard physiological properties.

### Running modulations are large in the peripheral representation in marmoset V1

We then tested whether V1 responses changed systematically during running. As in our prior report [**liska**], marmosets viewed a sequence of full-field drifting gratings, which elicited strong responses as expected.

We first assessed whether the time course of ensemble activity on V1 was correlated with the timeseries of running speeds. Consistent with our previous report, fluctuations in ensemble activity in foveal V1 did not appreciably correlated with whether the animal was running or not (Figure 2A). In striking contrast, simple visual inspection revealed that activity in peripheral V1 had clear increases in activity during times when the animal was running (Figure 2B). This is a clear difference between our foveal and peripheral datasets, despite their being recorded simultaneously.

**Fig. 2:**
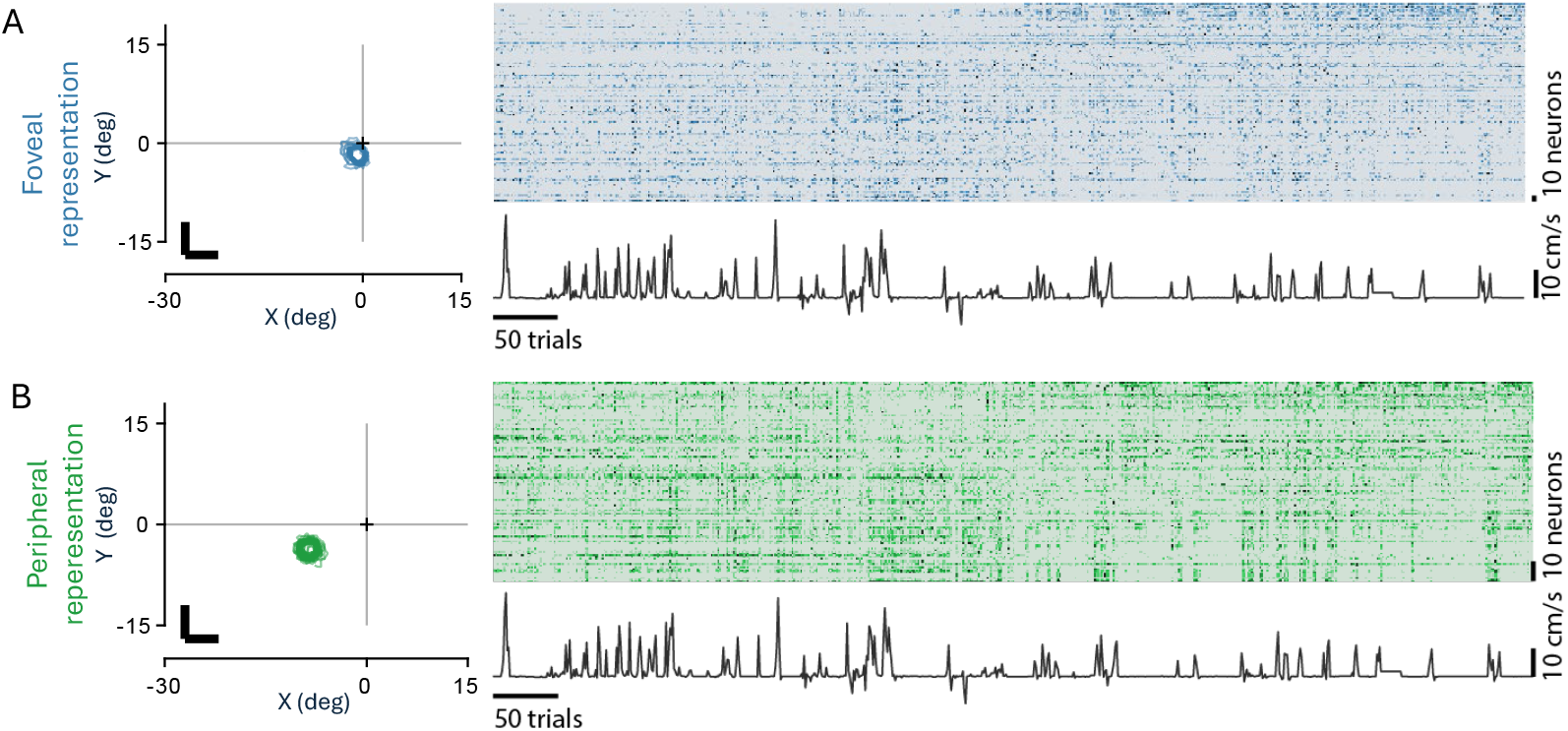
Example session for running with simultaneous recording. A. Left: Contour plots of receptive fields of neurons recorded from the proximal portion of the probe (the dorsal portion of V1), confirming foveal representations. Right, enseemble raster plot of spiking activity for this bank of neurons sorted in order of correlation with running speed (trace shown below). Correlation with running is not visually compelling. B. Same thing for deeper bank of V1 at calcarine sulcus, with peripheral representations. A correlation is visually evident. Note that the running trace is the same in both cases, as the foveal and peripheral datasets shown in this illustrative figure were recorded simultaneously.

We then performed a more quantitative unit-by-unit analysis, comparing the response of each unit to matched stimuli during periods of running versus not running. This also revealed a substantial increase in peripheral V1 activity during running, distinct from minimal effects in simultaneously-recorded foveal V1. Figure 3A shows the visually-driven response of each foveal V1 neuron averaged over all stimuli (i.e., over the entire orientation tuning curve), parsed by whether the animal was running (y-axis) or not running (x-axis). The resulting cloud of points is distributed fairly evenly about the line of unity, indicating no systematic effect of running. Figure 3B shows the same analysis for V1 neurons representing more peripheral portions of the visual field; that resulting cloud of points has a compelling amount of density above the line of unity, indicating a clear increase in activity during running. Quantitatively, these results correspond to a 20% increase in activity during running in the periphery and a 0.1% change in the fovea (Figure 3C).

**Fig. 3:**
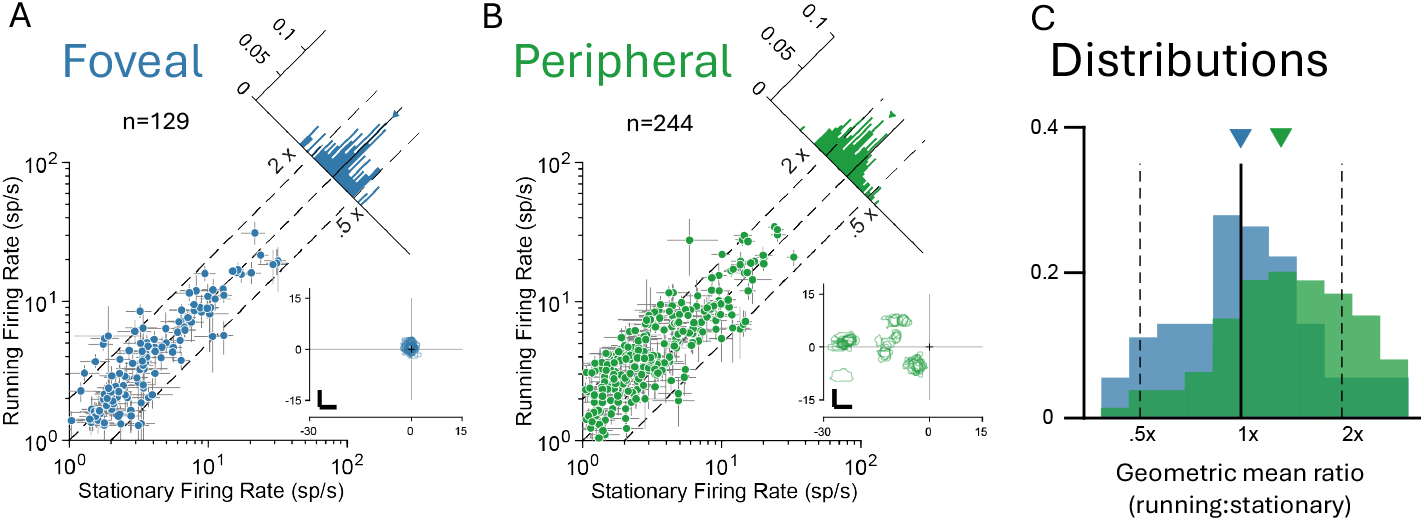
Running modulations of foveal and peripheral V1 summarized over units and combined across two animals. A. Scatterplot (log-log) for 129 neurons in foveal V1, with mean firing rate during stationary periods represented by the x coordinate and mean firing rate during running periods represented by the y coordinate with 95% confidence in grey. Histogram summarizes the projections onto the line of unity with a negligible shift during running (geometric mean ratio 1.01). Inset shows RF locations. B. Same for 244 neurons with peripheral representations. The histogram shows a significant shift indicating increases in response during running (geometric mean ratio 1.33). C. Comparison of distributions, note histograms now show higher gain on the right.

## Discussion

In our preceding report [3], we focused on a species difference in the magnitude (and sign) of running-correlated modulations in mouse and marmoset. The main result there was that V1 activity was barely different when the marmoset was running as compared to when they were stationary. But in that initial report, we noted the possibility of a difference in running-correlated modulations in the peripheral representation. Here, we report analyses of more comprehensive measurements that demonstrated a substantial increase in activity during running.

These results have implications both for the apparent species difference between rodents and primates in how (or whether) movement affects activity in V1, as well as understanding of how the cortical processing of information from the fovea (i.e, representing the central visual field) might differ from that of peripheral processing. Initial interpretations of our first report may have simplified the apparent species difference, reducing things down to a simple story that running has big effects on mouse V1 but has little if any effect on monkey V1. Our new results reported here condition that interpretation in an important way: the peripheral representation of V1 appears to be substantially affected by running. Although the size of the effect in peripheral V1 (30%) is considerably smaller than the effects seen in mouse V1, such a modulation is large relative to, say, attentional effects typically seen in primate V1 [**talluri**]. These modulations should therefore be considered substantive, as a change of activity on order of one-quarter to one-third of the stimulus-driven activity.

This substantial increase in activity of peripheral V1 activity, in light of the little-to-no effects in the fovea (perhaps, a very small reduction) also suggest that the primate foveal representation may be relatively unique. Much more data are needed to comparatively study both basic visual processing as well as effects of nonvisual study across the visual field, but for now our results invite intriguing speculation. We hypothesize that the cortical representation of the fovea (at least within V1) is itself specialized, following the specializations of the actual fovea in the retina. It is appealing to think that the fovea is relatively immune from behavioral modulations in order to allow for unambiguously image-based information to be processed at high resolution, whereby the peripheral representation might carry signals that are better modulated by behavioral context earlier. The validity of this hypothesis, and how it plays out in later stages of visual processing, now motivates further work and an integration of fine-grained anatomical and functional studies that will be possible in marmosets. For example, anatomical differences in feedback circuits across the visual field representation in V1 [9] are deserving of further consideration in light of our physiological findings.

## Materials and methods

We performed electrophysiological recordings in V1 of two common marmosets (1 male, “marmoset G”, and 1 female, “marmoset B”). Recordings were performed using Neuropixels 1.0 probes [10] acutely inserted into small craniotomies (procedure described below) over V1 in right hemisphere. All experimental protocols were approved by The University of Texas Institutional Animal Care and Use Committee and in accordance with National Institute of Health standards for care and use of laboratory animals.

Subjects stood quadrupedally on a 12” diameter wheel while head-fixed facing a 24” LCD (BenQ) monitor (resolution = 1920×1080 pixels, refresh rate = 120 Hz) corrected to have a linear gamma function, at a distance of 36 cm (pixels per degree = 26.03) in a dark room. Eye position was recorded via an Eyelink 1000 eye tracker (SR Research) sampling at 1 kHz. A syringe pump-operated reward line was used to deliver liquid reward to the subject. Timing events were generated using a Datapixx I/O box (VPixx) for precise temporal registration. All of these systems were integrated in and controlled by MarmoView. Stimuli were generated using MarmoView, custom code based on the PLDAPS [11] system using Psychophysics Toolbox [12] in MATLAB (Mathworks). In electrophysiology data gathered using Neuropixels probes, data were sent through Neuropixels headstages to a Neuropixels PXIe acquisition card within a PXIe chassis (National Instruments). The PXIe chassis sent outputs to a dedicated computer running Open Ephys with an Open Ephys acquisition board additionally attached to record timing events sent from the Datapixx I/O box. Spike sorting was performed using Kilosort 2.5.

### Acute Neuropixels recordings

Acute Neuropixels recordings were performed using standard Neuropixels 1.0 electrodes (IMEC, Leuven, Belgium). Each probe consists of 384 recording channels that can individually be configured to record signals from 960 selectable sites along a 10 mm long, 70 × 24 µm cross-section straight shank. Probes were lowered into right dorsal V1 via burr holes spaced irregularly along the AP axis 4-5 mm from the midline for a single session of experiments. Natural images were played to provide visual stimulus as well as occupy the subject and keep them awake during insertion and probe settling. The temporary seal on the burr hole was removed, the intact dura nicked with a thin needle and the burr hole filled with saline. The probe was then lowered through the dural slit at 500 µm/minute, allowing 5 minutes for settling every 1000 µm of total insertion. The whole-probe LFP visualization was monitored during insertion for the characteristic banding of increased LFP amplitude that characterizes cortical tissue. The probe was inserted to approximately 5.5mm depth below dura such that a clear second LFP band was visible to ensure that electrodes covered the full depth of the calcarine cortex. The probe was then allowed to settle for 30 minutes. Active electrode sites on the probe were configured to subtend both dorsal and calcarine cortex simultaneously. Post-hoc receptive field recreation confirmed that visually-driven, tuned, V1 neurons were recorded at both foveal and peripheral eccentricities.

### General experimental procedure

Marmoset recording sessions began with eye tracking calibration. Once calibration was completed, the wheel was unlocked and the subject was allowed to locomote freely, head-fixed, while free-viewing stimuli. Trials for all stimuli were 20 sec long with a 500 ms ITI and a 20 sec long natural image interleaved every fifth trial to keep the subject engaged. Stimuli were shown in blocks of 10 minutes and a typical recording session consisted of 50 trials of calibration followed by 1 or 2 blocks of a drifting grating stimulus and 1 block each of the two mapping stimuli. To elicit sufficiently reliable and frequent running behavior, subjects were rewarded at set locomotion distance intervals unrelated to the stimulus or gaze behavior (typical rewards were 50-70 µL and distance required to achieve a reward usually varied between 20-75 cm; reward amounts and intervals were adjusted daily to maximally motivate the subject.)

### Eye tracking calibration

While the wheel was locked, subjects were allowed to free-view a sequence of patterns of marmoset faces. Marmosets naturally direct their gaze towards the faces of other marmosets when allowed to free-view with little-to-no training, allowing for the experimenter to adjust the calibration offset and gain manually between pattern presentations. Faces were 1.5 degrees in diameter and were presented for 3 sec with a 2 sec ISI between patterns. A portion of presented patterns were asymmetrical across both the X and Y axes of the screen to allow for disambiguation in the case of axis sign flips in the calibration. 50 trials were presented before each recording session to verify and refine the calibration. Calibration drift between sessions was minimal, requiring minor (<1 deg) adjustments over the course of 1-2 months of recordings.

### Drifting grating stimuli

The primary stimulus consisted of full-field drifting gratings. Gratings were optimized to drive marmoset V1 with 3 separate spatial frequencies (1, 2, and 4 cycles per degree), two drift speeds (1 or 2 degrees per second) and 12 orientations (evenly-spaced 30 degree intervals). Each trial consisted of multiple grating presentations, each with a randomized spatial frequency, drift speed, and orientation. Gratings were displayed for 833 ms followed by a 249-415 ms randomly jittered inter-stimulus interval. After each 20 second trial there was a longer 500 ms inter-trial interval. Every fifth trial was replaced with a natural image to keep subjects engaged and allow for visual assessment of calibration stability on the experimenter’s display.

### Mapping of receptive fields

A spatiotemporal receptive field mapping stimulus, consisting of sparse dot noise, was shown during each recording session. One hundred 1 degree white and black dots were presented at 50% contrast at random points on the screen. Dots had a lifetime of 2 frames (16.666 ms). Marmosets freely viewed the stimulus and we corrected for eye position offline to estimate the spatial receptive fields using forward correlation [8].

### Analysis of tuning

We counted spikes between the 50ms after grating onset and 50ms after grating offset and divided by the interval to generate a trial spike rate. To calculate orientation tuning curves, we computed the mean firing rate each orientation and spatial frequency. Because we were limited by the animal’s behavior to determine the number of trials in each condition (i.e., running or not), we computed orientation tuning as a weighted average across spatial frequencies with with weights set by the spatial frequency tuning. We used these resulting curves for the all analyses of tuning. We confirmed that the results did not change qualitatively if we either used only the best spatial frequency or marginalized across spatial frequency.

Orientation selectivity index was calculated using the following equation

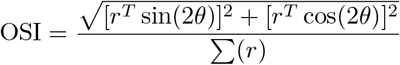

where *θ* is the orientation and *r* is the baseline-subtracted vector of rates across orientations.

## Acknowledgements

We thank Allison Laudano for animal and colony management and care, Christopher Badillo for apparatus design and fabrication, and Lily DeFelice for assistance with animal work.

## Data and materials availability

All data in the main text or the supplementary materials are available upon request, and will be posted publicly at time of publication.

